# Cell type-specific signal analysis in EWAS

**DOI:** 10.1101/2021.05.21.445209

**Authors:** Charles E. Breeze

**Affiliations:** Altius Institute for Biomedical Sciences, Seattle, WA, USA 98121; UCL Cancer Institute, University College London, London WC1E 6BT, UK

**Keywords:** DNA methylation, epigenome-wide association study, differentially methylated position, differentially variable position

## Abstract

Hundreds of epigenome-wide association studies (EWAS) have been performed, successfully identifying replicated epigenomic signals in processes such as ageing and smoking. Despite this progress, it remains a major challenge in EWAS to detect both cell type-specific and cell type confounding effects impacting study results. One way to identify these effects is through eFORGE (experimentally derived Functional element Overlap analysis of ReGions from EWAS), a published tool that uses 815 datasets from large-scale mapping studies to detect enriched tissues, cell types and genomic regions. Here, I show that eFORGE analysis can be extended to EWAS differentially variable positions (DVPs), identifying target cell types and tissues. In addition, I also show that eFORGE tissue-specific enrichment can be detected for sites below EWAS significance threshold. I develop on these and other analysis examples, extending our knowledge of eFORGE cell type- and tissue-specific enrichment results for different EWAS.

## 1. Introduction: Functional overlap analysis, FORGE and eFORGE

To give a background on functional element overlap analysis I will first introduce a method in this area that originated from research on genome-wide association studies (GWAS). GWAS are a key tool to uncover genetic associations with complex traits and diseases. During the last two decades, thousands of GWAS have been performed, and many thousands of single nucleotide polymorphisms (SNPs) have been associated with a range of phenotypes [1]. One critical component in the study of genetic associations with complex traits or diseases is interpreting the mechanism of action of identified variants. Many GWAS SNPs lie in non-protein-coding regions of the genome [2], and the lack of information regarding these regions has proved an obstacle for GWAS interpretation. This obstacle is in part addressed by large-scale efforts to chart regulatory elements genome-wide across a range of tissues and cell types, such as the ENCODE [3], Roadmap Epigenomics [4], and BLUEPRINT projects [5]. Intersection of these epigenetic maps, notably chromatin accessibility profiles (such as those obtained by DNase-seq), with disease-associated SNPs can help to pinpoint key cell types and tissues involved in disease mechanisms [2]. This approach can also help identify specific regulatory elements and downstream mechanisms involved [6].

In order to automate the complex process of identifying key cell types and tissues associated with a given GWAS, in 2014 Dunham et al. developed the FORGE (Functional element Overlap analysis of the Results of GWAS Experiments) tool [7]. The FORGE tool uses a method termed Functional element Overlap Analysis (FOA) to identify key cell types and tissues associated with a list of top GWAS SNPs from a particular study.

In order to verify whether a list of SNPs is enriched in epigenetic mark peaks from a given tissue sample, FORGE generates background lists of SNPs to compare to the input list in terms of overlap with epigenetic mark peaks. These background lists are designed to contain SNPs similar to SNPs from the input list for a range of genomic annotations. For example, if 50% of the SNPs in the input list overlap a transcription start site (TSS), then 50% of the SNPs in each of the background lists will also overlap a TSS. In fact, there are three different levels of annotation that are considered when ensuring similarity between background SNP lists and the input SNP list. These three levels are: distance to TSS, minor allele frequency (MAF) and guanine-cytosine (GC) content. For each of these three annotation levels, different groups or bins are created to reflect the varying level of distance to TSS, MAF, or GC content for different sets of SNPs. For example, one bin will contain SNPs that directly overlap (or are very close to) a TSS, while another bin will contain SNPs that are further away from the TSS up to a specified distance, yet another bin will contain SNPs that are even further, and so on, to complete the annotation of TSS distance for every SNP in the background pool.

For a SNP to be in the background pool of SNPs in FORGE, it must be contained in one of the following arrays: Illumina_HumanOmni2.5 array, Illumina_Human1M-duoV3, Illumina_Human660W-quad, HumanHap300v2, HumanHap550v3.0, Illumina_Cardio_Metabo,Affy_GeneChip_100K_Array, Affy_GeneChip_500K_Array, Affy_SNP6, HumanCNV370-Quadv3 [7].

FORGE also considers linkage disequilibrium (LD) for analysis. Specifically, FORGE filters SNPs that are in the input list of SNPs for LD, selecting only one SNP from each LD block (r squared>0.8), to avoid including multiple proximal SNPs that are representative of one single genomic association and that may even overlap the same epigenetic mark peak.

FORGE analysis across GWAS using DNase-seq hotspot data from the ENCODE and Roadmap Epigenomics consortia uncovered a range of tissue-specific associations with complex traits and diseases [7]. Given that FORGE performs many tests for enrichment, a multiple testing correction procedure (specifically Bonferroni testing at tissue level) was included in the FORGE code. In addition, as a particular disease-associated GWAS SNP list typically may contain a higher number of SNPs overlapping DNase-seq hotspots in a key tissue for the relevant disease, FORGE also reports the specific SNPs that overlap DNase-seq hotspots in each sample. These SNPs are reported in static and interactive tables, for further study and functional characterisation.

Thus, FORGE is a comprehensive tool for identifying key tissues, epigenetic datasets, and specific regulatory elements associated with SNPs from a given GWAS. Given the success of the Functional element Overlap Analysis (FOA) approach in GWAS, in 2016 Breeze et al. decided to implement this approach for EWAS performed with Illumina 450k and 850k arrays [8]. The resulting method was termed eFORGE (experimentally derived Functional element Overlap analysis of ReGions from EWAS).

## 2. Materials

eFORGE is available through a web browser (https://eforge.altiusinstitute.org/). Standalone eFORGE has been successfully installed and run both in cluster (Red Hat Linux, CentOS 7) and laptop environments (OS X 10.9.5) [8]. eFORGE code is available from https://github.com/charlesbreeze/eFORGE. eFORGE databases can be downloaded from https://eforge.altiusinstitute.org/?download.

## 3. Methods

### 3.1 Functional overlap analysis and EWAS: The design of eFORGE

Given the differences between GWAS and EWAS, including differences in the annotation of SNPs and DNAm positions, eFORGE presents some important changes with respect to FORGE.

Specifically, eFORGE does not apply SNP-centric metrics such as MAF or LD for the analysis of differentially methylated positions (DMPs) from a given EWAS. Instead, eFORGE selects background 450k/850k probes based on gene-centric categories (1st Exon, 3’ untranslated region or UTR, 5’UTR, Body, intergenic region or IGR, TSS1500 and TSS200) and CpG island-centric categories (CpG island, CpG island shore/shelf, NA or “open sea”). For example, if 70% of probes in the input list from an EWAS are in a CpG island, then 70% of probes in each of the background probe lists will be in a CpG island. If 5% of the input list probes are located in intergenic regions and outside of CpG islands, shelves, and shores, then 5% of the probes in the background lists will be in intergenic regions and outside of CpG islands, shelves, and shores. Thus, background annotation is different in eFORGE when compared to FORGE.

In addition, eFORGE does not filter input probes for LD but instead uses a 1kb proximity filter, as DNA methylation (DNAm) levels have been shown to correlate highly within 1kb [9]. This way eFORGE avoids testing the same DNA methylation signal twice in the analysis of DMPs from EWAS.

Finally, eFORGE performs a different approach related to accounting for the multiple tests involved during analysis runs. Instead of using the Bonferroni method, eFORGE applies a false discovery rate (FDR) approach, specifically the Benjamini-Yekutieli (BY) approach, which accounts for the non-independence between datasets [10]. Because eFORGE performs a cell-type level BY correction, eFORGE can identify enrichments at a cell type level, and not just a general tissue level (which is the case for FORGE analysis).

### 3.2 Cell type-specific and cell type-confounding effects in eFORGE

eFORGE was designed to test whether disease-associated DMPs from EWAS present increased overlap with regulatory elements (and thus epigenetic mark peaks) for tissues with an involvement in the disease under study. Cell type-specific effects in EWAS merit special attention, as DNAm differences due to cell type proportions (termed cell composition effects) are an important source of confounding. Cell composition effects are present when the proportion of a given cell type in the sampled tissue varies between EWAS cases and controls. For example, there may be a higher proportion of CD4+ T cells in whole blood in cases when compared to controls. If this occurs, then this is a case of confounding, and a lot of apparently phenotype-driven DMPs will actually be CD4+ T cell-specific DNAm positions. In this case, there is not really a change in the DNAm status of a particular genomic position due to disease (the goal of many EWAS), but rather a change in cell proportions. This problem of cell composition effects is one of the main problems affecting the field of EWAS.

One important distinction to be made with regard to cell proportions and EWAS DMPs is the distinction between cell composition effects and cell type-specific effects. Cell composition effects, as mentioned before, are important confounders in EWAS and represent regions of cell type-specific stable DNAm. On the other hand, some DNAm changes observed in EWAS may be true dynamic changes within particular cells that are phenotype-driven and occur in only a subset of the cell types assayed in each sample. For example, a true DNAm change may only occur within CD4+ T cells in a whole blood-based EWAS. When true, non-confounding DNAm changes occur in a subset of cell types from an EWAS tissue sample, these are termed cell type-specific effects. Unlike cell composition effects, cell type-specific effects represent true findings in EWAS research.

FOA as implemented in eFORGE can identify both cell type-specific effects and cell composition effects in EWAS. Typically, for a particular analysis on a whole blood-based EWAS, a very strong enrichment skewed towards a specific cell type in eFORGE can be indicative of cell composition effects, while analysis on cell composition corrected studies typically shows a weaker (yet significant) enrichment as characteristic of cell type-specific effects. For example, in the analysis of the cell composition corrected EWAS on rheumatoid arthritis (RA) by Liu et al., eFORGE shows an enrichment for immune cells [8],[11]. On the other hand, eFORGE analysis of non-cell composition corrected whole blood EWAS can uncover strong enrichments skewed towards a specific dataset, identifying the key cell types and tissues involved in both contexts [8]. Thus, eFORGE can be used to identify both cell type-specific effects and cell composition effects in EWAS.

Interestingly, eFORGE can also be used to test notions of tissue mimicry in EWAS. Some researchers suggest that epigenomic perturbations found in the target tissue can be reflected or shared in DNAm changes in surrogate tissues [12]. Thus, in this context tissue mimicry is suggested to occur if DNAm changes in one sampled tissue (e.g. blood) were representing or “mimicking” DNAm changes in a nonsampled tissue (e.g. ovary), indicating shared effects in both tissues. An example of tissue mimicry or shared effects across tissues is the smoking-induced differential methylation of the AHRR gene across blood, skin, adipose and lung tissue [13]. One specific case where tissue mimicry was tested with eFORGE was for a blood DNAm signature predictive of ovarian cancer status [14]. eFORGE analysis of the top DMPs from this study, however, shows an blood-specific regulatory element enrichment that suggests that these sites are representative of an immune response signature to ovarian cancer rather than sites that are likely to be shared by DNAm changes in the ovarian cancer tissue itself [8]. Therefore, eFORGE can be used to aid the biological interpretation of EWAS results in the area of surrogate tissues and tissue mimicry.

eFORGE can also be applied to the study of differentially variable positions (DVPs) in EWAS. DVP analysis represents a very different approach to the standard DMP analysis, as the variance, instead of the mean, is analysed to obtain disease-associated positions [15]. eFORGE analysis on top DVPs from a published study on inter-individual DNAm variability in sorted blood cells [16] uncovers key cell types and regulatory regions involved, including a cell type-specific enrichment for monocytes and T cells (**Figure 1A, B**). Thus, eFORGE can also be applied to the study of differentially variable positions in EWAS.

**Figure 1:**
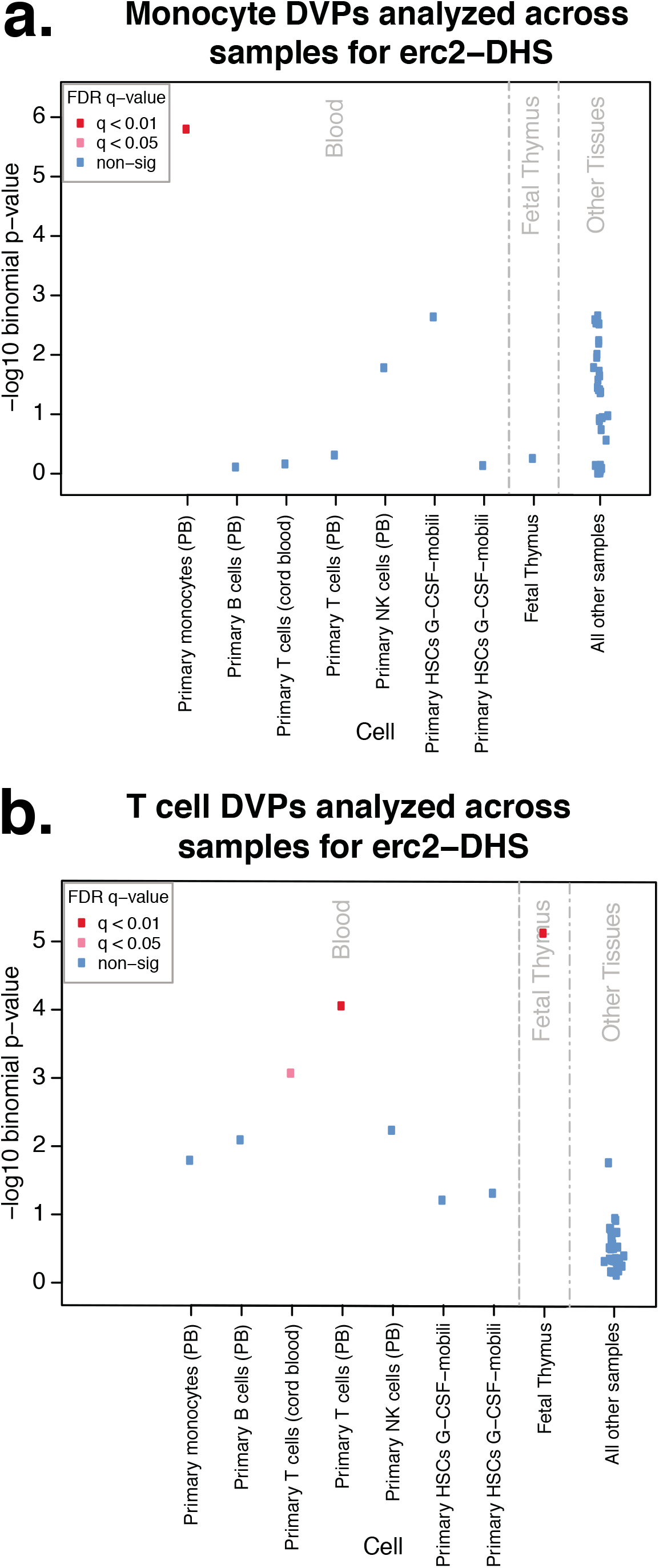
eFORGE analysis of inter-individual variation DVPs: **[A]** Analysis of 14 top Monocyte DVPs shows high monocyte-specific eFORGE DNase-seq hotspot enrichment (red dot, q-value <0.01). No other tissue or cell type is enriched (blue dots, 0.05 < q-value), indicating a unique signature of enrichment for monocyte DVPs in eFORGE DNase-seq hotspot analysis. Results indicate a highly significant overlap between monocyte inter-individual DVPs and monocyte DNase-seq hotspots from the Roadmap Epigenomics Consortium. PB: Peripheral Blood. HSC: Haematopoietic Stem Cell. **[B]** Analysis of 47 top T cell DVPs shows high T cell- and Thymus-specific eFORGE DNase-seq hotspot enrichment (red dots, q-value <0.01, and pink dot, q-value <0.05). No other tissue or cell type is enriched (blue dots, 0.05 < q-value), indicating a unique signature of enrichment for T cell DVPs in eFORGE DNase-seq hotspot analysis. Results indicate a highly significant overlap between T cell inter-individual DVPs and T cell DNase-seq hotspots from the Roadmap Epigenomics Consortium. PB: Peripheral Blood. HSC: Haematopoietic Stem Cell.

## 4. Specific notes on eFORGE analysis

When analysing EWAS data with eFORGE it is important to highlight several aspects:

### 4.1 Number of probes

eFORGE analysis builds a distribution of background peak overlaps and then compares this distribution to the peak overlap number for the input probe list. This way, eFORGE can compute a binomial p-value. In order for the background distribution to be truly representative, there has to be an appropriate number of probes in the input list (over 5 -but preferably over 100-probes). This way, it is possible to ensure an adequate number of probes in the background probe lists. However, achieving a sufficiently high number of probes is a problem for some EWAS. For example, certain studies have only one or two significant probes that are associated with a given trait (significant at a p-value <10-7 or some appropriate threshold to account for multiple testing). In this context, it can be shown that eFORGE tissue-associated enrichment can continue below a defined p-value significance threshold (**Figure 2**), much in the same way as tissue-associated enrichment can be detected below genome-wide significance in GWAS [2]. Because of this, it is recommended that users test top 100 and top 1000 sites even if they may be below the nominal significance threshold in EWAS, as tissue-specific signal in cases may still hold below this threshold. This type of approach has also been shown to reveal further functional information, having been applied extensively in GWAS [2][17]. Exploring the dataset by testing a high number of top probes is recommended, especially up to 1000 probes (the input probe list limit in eFORGE web, which was set in consideration of gene-centric and CpG island-centric probe bin size limits used for background probe selection).

**Figure 2:**
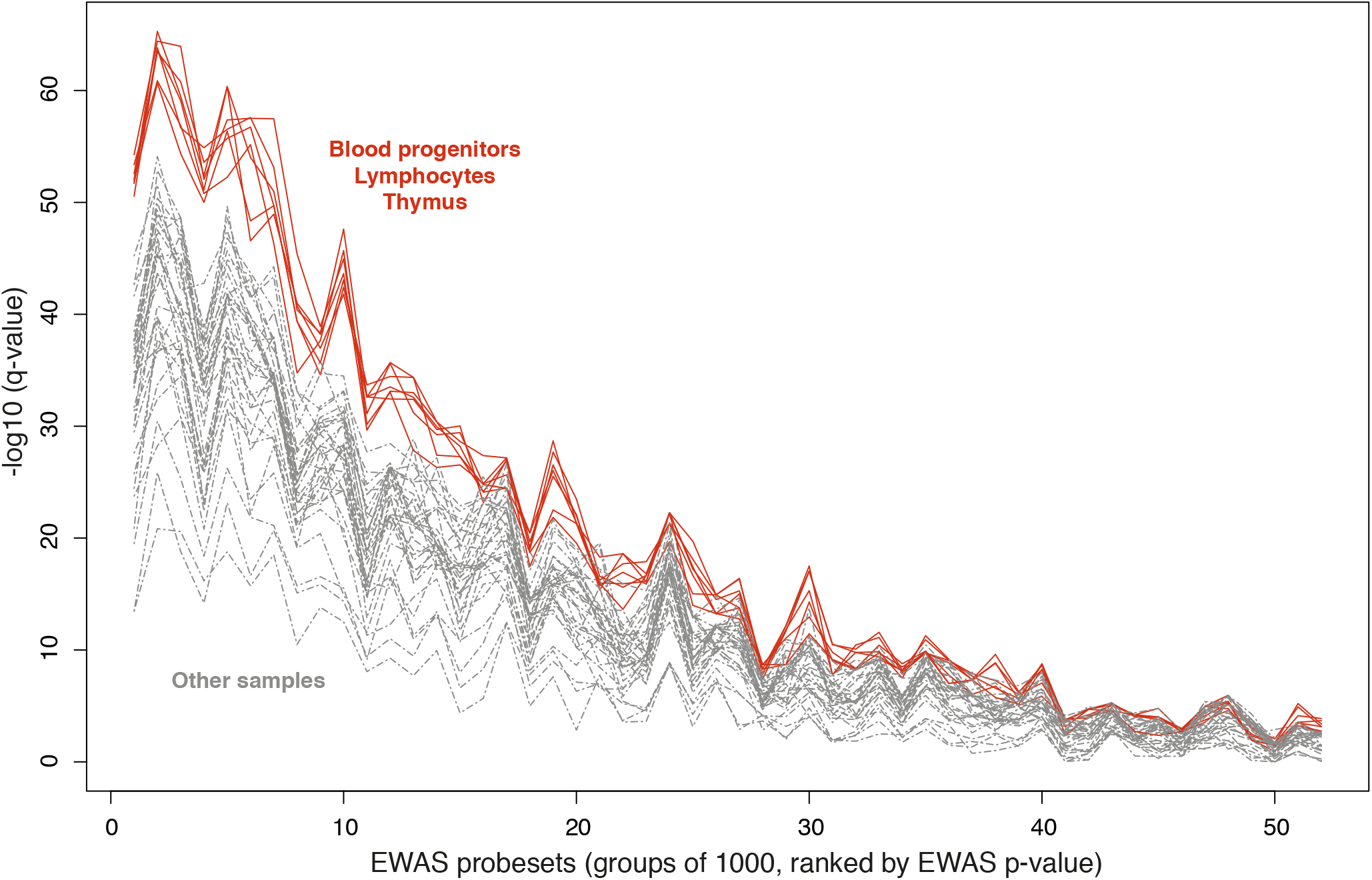
eFORGE analysis of RA-associated DMPs: −log10(q-values) were obtained from 52 eFORGE runs, each testing a different list of 1000 probes from the Liu et al. EWAS on RA, with probe lists ranked by EWAS p-value (lower p-value sets on the left side) [11]. A set of blood progenitors, lymphocytes and thymus samples (red) is detected as preferentially enriched when compared to a mixed set including all other samples (grey). Each eFORGE run was performed analysing the 2015 Roadmap Epigenomics DNase-seq hotspot dataset with 1000 background repetitions and other default settings.

### 4.2 Separating hypo- and hypermethylated sites

EWAS DMPs segregate into two different categories, hypo- and hypermethylated positions. Hypomethylated positions for a given disease or phenotype are those that present lower DNA methylation values in cases when compared to controls. Conversely, hypermethylated positions are those the present higher DNA methylation levels in cases when compared to controls. Of course, these DNA methylation levels are representative of a collection of cells in a tissue, as for each individual allele in each cell DNAm is either present or absent. It is at the cell population level where this gradual metric of DNAm is quantified.

In eFORGE analysis of EWAS DMPs for a particular study, it is often useful to separate hypo- and hypermethylated DMPs to obtain further insights into cell type- and tissue-specific associations. For example, eFORGE analysis of the top 1000 hypermethylated probes from a study on pre-invasive lung cancer lesions [18] reveals a strong enrichment for stem cell-specific DNase-seq hotspots (Roadmap Epigenomics 2015 data, **Figure 3A**). Conversely, analysis of the top 1000 hypomethylated probes from the same EWAS reveals no enrichment for any tissue or cell type (**Figure 3B**). In addition, analysis of a combined set of the top 500 hypomethylated and the top 500 hypermethylated probes (1000 probes in total) also reveals no enrichment for any tissue or cell type (**Figure 3C**). In view of these results, and given the different regulatory effects of hypo- and hypermethylated DMPs, it is important to at least consider separating hypo- and hypermethylated top probes when analysing EWAS data with eFORGE.

**Figure 3:**
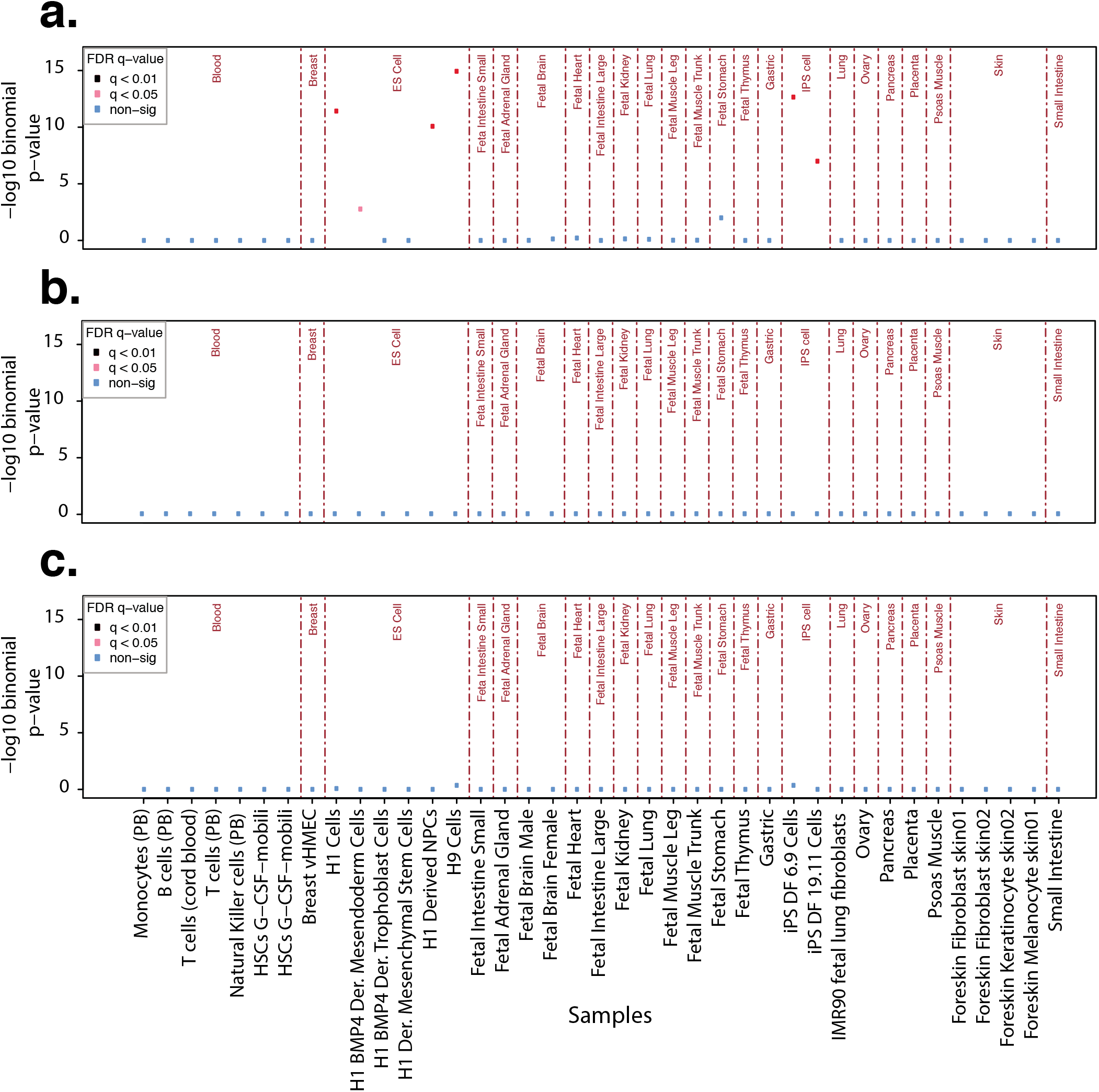
eFORGE analysis of DMPs associated with progressive pre-invasive lung cancer lesions: **[A]** −log10(q-values) obtained from eFORGE runs on the top 1000 hypermethylated positions from the lung cancer lesion EWAS by Teixeira et al. [18], analysis performed using the Roadmap Epigenomics 2015 DNase-seq hotspot dataset, with 1000 background repetitions and other default settings. **[B]** Same for the top 1000 hypomethylated positions. **[C]** Same but for a combined set of the top 500 hypomethylated positions and the top 500 hypermethylated positions (1000 probes in total). HSC: Haematopoietic Stem Cells. NPC: Neuronal Progenitor Cells. PB: Peripheral Blood. Der.: Derived.

### 4.3 Tissue-specific, general and mixed signals

There are three classes of significant eFORGE results that have been observed in the analysis of EWAS: tissue-specific, general, and mixed signals.

#### 4.3.1 Tissue-specific eFORGE results

these results are observed when EWAS DMPs significantly co-localise with regulatory elements that are particular to a specific tissue. This class of results is quite common, for example, analysis of three EWAS on autoimmune diseases (SLE, Sjögren’s syndrome, and RA) uncovers a significant enrichment for immune cells and related immune tissues for all three traits [8]. Given that the SLE and Sjögren’s syndrome EWAS were performed on naïve CD4+ T cells, and the RA EWAS underwent cell composition correction, these associations point to true cell type-specific effects within blood, associated with changes in immune regulatory elements and programs. For EWAS where cell composition correction has not been applied, disproportionately strong tissue-specific results may be indicative of confounding cell composition effects. In addition, tissue- (or cell type-) specific eFORGE enrichments can also be observed when testing tissue-specific differentially methylated positions (tDMPs) and cell type-specific differentially methylated position (ctDMPs).

#### 4.3.2 General enrichment results

these results are observed when EWAS DMP sets significantly co-localise with epigenetic mark peaks across all tissues. This may occur, for example, if they affect regulatory elements controlling housekeeping genes or similar shared regulatory processes. General enrichment results have been observed in published EWAS [19], and represent a distinct class of eFORGE results, separate from the absence of eFORGE signal, and separate from both tissue-specific eFORGE results and mixed enrichment eFORGE results.

#### 4.3.3 Mixed enrichment

sometimes eFORGE results do not clearly point to a particular tissue, but reveal enrichment for several (yet not all) tissues. These enrichments represent a significant co-localisation with regulatory elements shared across a specific group of tissues. This can typically occur in related tissues, and in these cases may be representative of shared lineage-specific regulatory programs. Mixed enrichment represents a distinct class of eFORGE enrichments that merits further investigation.

### 4.4 Choosing different datasets to analyse in eFORGE

Several different epigenetic track datasets are available for eFORGE analysis: DNase I hotspots from the Roadmap consortium (2012 and 2015 releases), histone mark broadPeaks (Roadmap consortium), Hidden Markov Model (HMM) chromatin states (Roadmap consortium), and DNase I hotspots from the ENCODE and BLUEPRINT consortia. These analysis options segregate into 3 broad epigenetic track classes: accessible chromatin tracks, histone mark broadPeak tracks, and HMM chromatin state tracks.

The accessible chromatin track represents accessible regions throughout the genome, indicative of regulatory elements like enhancers, insulators, and promoters. This class also includes structural elements such as CTCF binding sites. Enrichment in an accessible chromatin track for a particular cell type suggests EWAS sites are associated with genomic regulation in that cell type. In terms of association with mechanisms of gene regulation, accessible chromatin is one of the broadest analysis categories that can be assayed (for example through DNase-seq and ATAC-seq).

On the other hand, histone mark broadPeak enrichments may point to an association with particular classes of regulatory elements and processes. For example, some histone marks are associated with active genes (e.g. promoter-associated H3K4me3) and other histone marks are associated with repressed genes (e.g. polycomb-associated H3K27me3), and therefore specific eFORGE histone mark enrichments may indicate an association with regions belonging to sets of active or repressed genes in a particular tissue. Further subdivision in cell type enrichment signal can be found for different levels of active histone mark broadPeaks. For example, marks associated with gene activation segregate into the promoter-associated H3K4me3, the enhancer-associated H3K4me1, and the transcription-associated H3K36me3. Conversely, histone marks associated with gene repression segregate into the polycomb repressed region-associated H3K27me3 and the heterochromatin-associated H3K9me3. In order to view enrichments across all these regulatory element classes, it is recommended that all histone mark broadPeak tracks be analysed in parallel (“H3 all” option), so that the relative enrichment across different histone marks can be compared.

Finally, HMM chromatin states tracks are similar to histone mark tracks in that different chromatin states are also representative of different classes of active and repressed regulatory elements. In fact, given the high number of chromatin states (15), eFORGE always analyses all chromatin states in parallel (“chromatin state” option), so that enrichments across chromatin states can be compared on an equal basis.

Histone mark broadPeaks and HMM chromatin states are especially valuable for eFORGE analysis as they provide a deeper insight into the potential regulatory associations of the identified regions, for example, by highlighting associated enhancers and promoters. Probe subsets derived from these enhancer- and promoter-associated lists can allow for the construction of more detailed tissue-specific regulatory annotations of EWAS results and have the potential to aid downstream pathway analysis.

### 4.5 False positive rate

eFORGE test runs on randomly selected 450k and 850k datasets of different sizes reveal a remarkably low level of false positives. On average, close to 7 in 100,000 tests result in a significant association (q-value<0.01). Testing the recommended number of probes (e.g. 100 DMPs) results in a level of false positives below 1 in 100,000 tests, which is lower than for other comparable tools [7], confirming previous results [8].

### 4.6 Reproducibility of eFORGE results

Given that eFORGE randomly samples the background set of probes each time the tool is run, a repeated set of eFORGE analyses may yield slightly different q-values (due to differences in the overlap counts for randomly selected probe lists in each run). This variability is low, especially for the default case of 1000 background repetitions and testing a list with over 100 probes. In these cases, eFORGE q-values have been shown to vary with a standard deviation of 0.011 for an example set of probes from the Altorok et al. EWAS on Sjögren’s syndrome [20] (**Figure 4**). This low level of variability indicates that eFORGE analyses are reproducible, confirming previous results [8].

**Figure 4:**
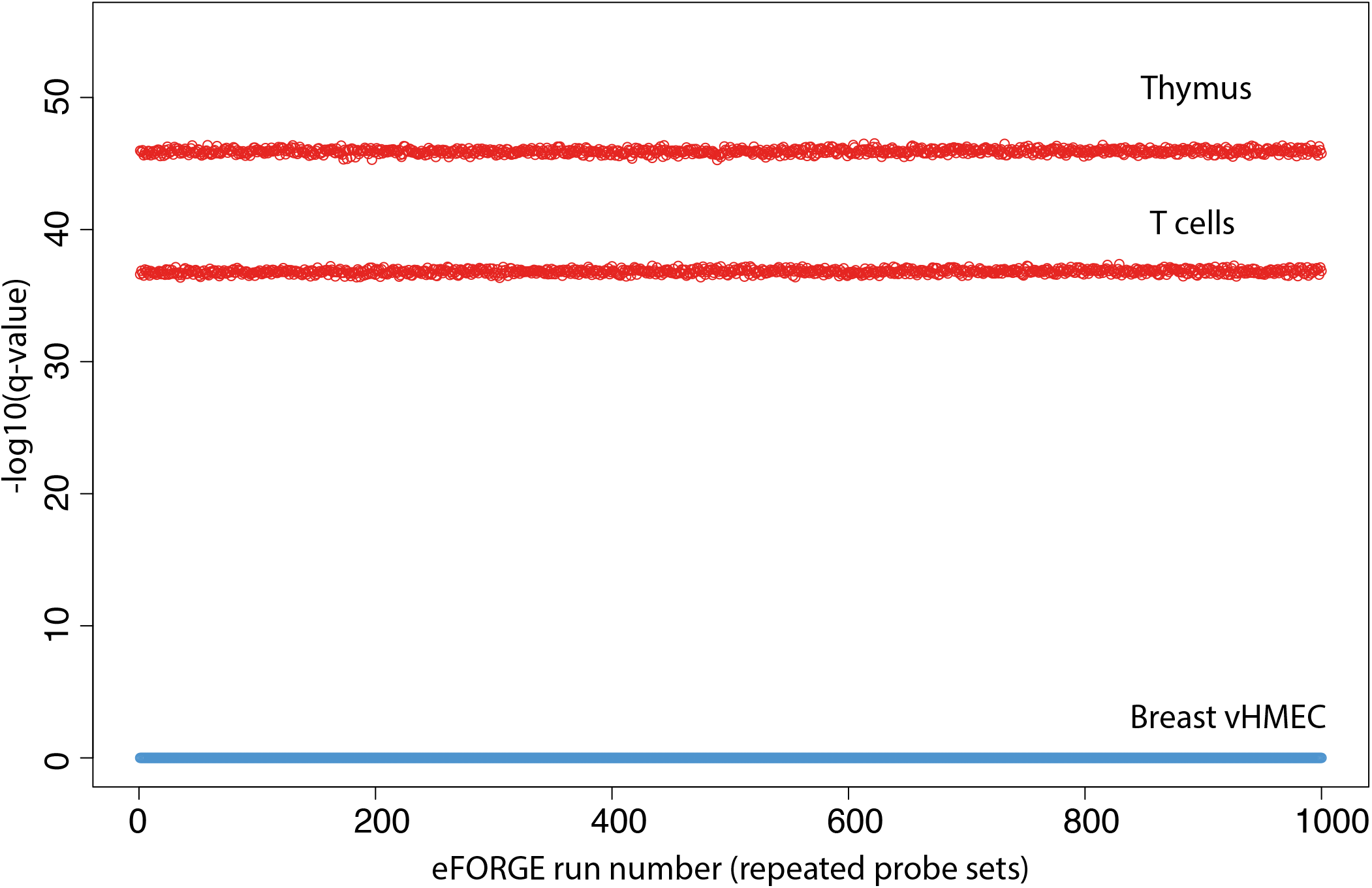
repeated eFORGE analyses of Sjögren’s syndrome-associated DMPs: −log10(q-values) were obtained from 1000 distinct eFORGE runs, each performed with 1000 background repetitions testing a list of 753 top probes from the Altorok et al. EWAS on Sjögren’s syndrome [20]. eFORGE analysis was performed using the 2012 Roadmap Epigenomics DNase-seq hotspot dataset, with default settings. Both T cell- and Thymus-specific enrichments are reproducibly detected across the 1000 runs. Non-enriched values for Breast vHMEC (variant Human Mammary Epithelial Cells) are also shown across the 1000 runs.

### 4.7 Evaluating enrichments across replicate samples

When analysing EWAS results in eFORGE “erc 2012” mode, analysis is performed across 299 samples, many of which are replicates within a given tissue. This analysis provides an insight into eFORGE results across multiple samples of the same tissue. Despite the fact that some of the samples may present higher or lower quality in the mapping of accessible chromatin sites, all samples included were of sufficient quality to be present in the Roadmap Epigenomics 2012 data release. If an eFORGE enrichment is observed for many replicate samples of a given tissue, then this provides further evidence for the presence of a tissue-specific enrichment. However, if eFORGE results are only observed for a small subset of the replicate samples of a given tissue, then caution in interpreting results is advised as it is important to consider the inherent variability in epigenetic mapping experiments. In these cases, further examination of the epigenetic track peaks and accessible regions underlying enrichment is advised.

### 4.8 Number of background repetitions

The number of background repetitions is set by default to 1000. Lower numbers are possible and will increase the speed of the tool. However, it is recommended to set the number of repetitions to 1000 to obtain a sufficiently large set of random background samples.

### 4.9 Enrichment or depletion

Apart from the standard eFORGE enrichment analysis, testing for the contrary of enrichment, depletion, is also available in eFORGE. A depletion test will show whether a set of EWAS probes tends to significantly not overlap a set of epigenetic mark peaks when compared to the background.

### 4.10 Significance threshold

Another option in eFORGE is to change the BY-corrected q-value significance threshold. Standard eFORGE analysis sets significant overlaps as those with a q-value < 0.01, with an intermediate level of significance at q-values < 0.05. It is not recommended that these values be changed from the default settings. However, for those seeking more stringent significance thresholds, those values are kept open as options.

### 4.11 Analysis label

During eFORGE analysis, a label can be provided. This label will be added to the static and interactive charts and tables that are generated by the tool, allowing users to keep track of the specific analyses run and record their own catalogue of eFORGE results.

### 4.12 Broad view and future directions

In order to obtain a broad view of eFORGE results across a variety of EWAS, a previously published effort selected studies with sample size over 100 from a review on published EWAS, and eFORGE analysis was performed across top probes from each study [8]. The results showed that tissue-specific eFORGE enrichments are frequent across EWAS, with 14 out of 20 enriched studies showing eFORGE tissue-specific enrichment [8]. The resulting 6 out of 20 studies presented mixed enrichment of two or more tissues, meriting further investigation [8]. Thus, this analysis demonstrated potential for eFORGE application across studies in the search of functional interpretation of EWAS results.

Several further steps have recently been taken to improve and extend eFORGE analyses of EWAS regions, including TF motif analysis and the design of a TF site-specific browser [21]. Given that TFs drive genomic regulatory programs and are associated with many of the epigenetic track peaks analysed by eFORGE, it is critical to further characterise the link between TF data and EWAS regions. Thus, a recent separate eFORGE module was created, termed eFORGE-TF, to provide further TF-related analysis [21].

First, a TF-centric browser was designed for viewing TF motif and footprint associations for over 850,000 probes [21]. This browser shows for each individual probe the overlap with TF motif matches (FIMO scan p-value<1e-04) and DNase-seq cutcount and footprint data (indicative of tissue-specific TF binding) across multiple tissues.

Second, a summary view was also provided to analyse a set of probes from a particular EWAS [21]. This analysis is aimed to detect TF motifs associated with a particular list of EWAS probes. In this view, for each associated motif, eFORGE-TF also provides the relative position of input EWAS probes to motif basepairs. This allows for further insights into the relationship between DNAm changes and TF motifs for a particular study.

Both of these resources, the browser and the summary view, can be accessed from eFORGE-TF at https://eforge-tf.altiusinstitute.org/ [21].

In summary, eFORGE represents a powerful resource for the analysis of EWAS DMPs, designed to detect EWAS-associated regulatory elements and TF binding sites, thus yielding further regulatory insights for the functional interpretation of EWAS results [8][21].

